# AT-hook DNA-binding motif-containing protein one knockdown downregulates EWS-FLI1 transcriptional activity in Ewing’s sarcoma cells

**DOI:** 10.1101/2022.05.16.492174

**Authors:** Takao Kitagawa, Daiki Kobayashi, Byron Baron, Hajime Okita, Tatsuo Miyamoto, Rie Takai, Durga Paudel, Tohru Ohta, Yoichi Asaoka, Masayuki Tokunaga, Koji Nakagawa, Makoto Furutani-Seiki, Norie Araki, Yasuhiro Kuramitsu, Masanobu Kobayashi

**Affiliations:** Advanced Research Promotion Center, Health Sciences University of Hokkaido, 1757, Kanazawa, Ishikari-Tobetsu, Hokkaido, 061-0293, Japan; Department of Omics and Systems Biology, Graduate School of Medical and Dental Sciences, Niigata University, 757 Ichibancho, Asahimachi-dori, Chuo-ku, Niigata, 951-8510, Japan; Department of Tumor Genetics and Biology, Faculty of Life Sciences, Kumamoto University, Kumamoto-Shi, Kumamoto, 860-8556, Japan; Center for Molecular Medicine and Biobanking, University of Malta, Msida, MSD2080, Malta; Division of Diagnostic Pathology, Keio University School of Medicine, Shinano, Shinjuku-ku, Tokyo, 160-8582, Japan; Department of Molecular and Cellular Physiology, Yamaguchi University Graduate School of Medicine, 1-1-1, Minami-kogushi, Ube, Yamaguchi, 755-8505, Japan; Department of Systems Biochemistry in Pathology and Regeneration, Yamaguchi University Graduate School of Medicine, 1-1-1, Minami-kogushi, Ube, Yamaguchi, 755-8505, Japan; Department of Obstetrics and Gynecology, Yamaguchi University Graduate School of Medicine, 1-1-1, Minami-kogushi, Ube, Yamaguchi, 755-8505, Japan

## Abstract

Ewing’s sarcoma is the second most common bone malignancy in children or young adults and is caused by an oncogenic transcription factor by a chromosomal translocation between the EWSR1 gene and the ETS transcription factor family. However, the transcriptional mechanism of EWS-ETS fusion proteins is still unclear. To identify the transcriptional complexes of EWS-ETS fusion transcription factors, we applied a proximal labeling system called BioID in Ewing’s sarcoma cells. We identified AHDC1 as a proximal protein of EWS-ETS fusion proteins. AHDC1 knockdown showed a reduced cell growth and transcriptional activity of EWS-FLI1. AHDC1 knockdown also reduced BRD4 and BRG1 protein levels, both known as interacting proteins of EWS-FLI1. In addition, AHDC1 co-localized with BRD4. Our results suggest that AHDC1 supports cell growth through EWS-FLI1.

## Introduction

Ewing’s sarcoma is the second most common bone malignancy in children or young adults. This tumor is caused by a chromosomal translocation of the EWS RNA binding protein 1 (EWSR1) and the E-twenty-six (ETS) transcription factor family, which mainly consists of the Friend Leukemia integration 1 (FLI1), ETS-related gene (ERG), E1A enhancer-binding protein (E1AF), or other kinds of ETS transcription factors [1, 2]. The EWS-FLI1 fusion protein, consisting of the EWSR1 gene and the FLI1 gene, which has a transcriptional activation site due to chromosomal translocation, is detected in more than 85% of cases in Ewing’s sarcoma.

Transcription factors have been undruggable because they do not have ligand-binding pockets that small molecules can recognize and do not have a folding structure [3]. Transcriptional complexes that interact with oncogenic transcription factors are promising targets but not direct inhibition for the oncogenic transcription factors. EWS-ETS fusion proteins need more co-operational transcription factors and co-transcriptional regulators for the oncogenic functions. Several interacting partners of EWS-ETS fusion proteins have been isolated as druggable targets [4]. RNA helicase A interacts with EWS-FLI1, and their interaction is inhibited by a small molecule, YK-4-279, resulting in reduced tumor growth in vitro and in vivo experiments [5]. PARP1 also interacts with EWS-FLI1, and PARP1 inhibitors inhibit tumor growth [6]. Recently, BRD4, one of the super-enhancers and a target of the BET inhibitor, also interacted with the EWS-ETS fusion protein and reduced tumor growth [7, 8]. Therefore, transcriptional complexes with the EWS-ETS fusion protein might be a druggable target.

The proximal protein biotinylation method has been developed to identify proximal complexes of the target proteins using the biotin identification (BioID) and the ascorbate peroxidase (APEX) method [9]. Roux *et al*. developed a BioID method that uses BirA mutant (R118G) to provide biotinyl-5’-AMP intermediate and induces non-specific biotinylation of the proximal proteins [10]. EWS-FLI1 interactome analysis using the BioID method has already been achieved in human embryonic kidney 293T (HEK293T) cells. This approach showed that the cation-independent mannose 6-phosphate receptor works as a transporter of lysosomal hydrolases via lysosome-dependent turnover of EWS-FLI1 [11].

The aim of this study is to identify new interacting proteins of EWS-ETS fusion proteins using BioID system in Ewing’s sarcoma cells and investigate whether these affects cell growth and transcription of EWS-ETS fusion proteins. Our approach identified AT-hook DNA-binding motif-containing protein 1 (AHDC1) as one of the proximal proteins for EWS-ETS fusion proteins. AHDC1 has been revealed as a responsible gene in Xia-Gibbs syndrome patients, which causes an autosomal dominant multisystem developmental disorder [12–17]. AHDC1 knockdown showed reduced protein levels of EWS-FLI1 or target proteins of EWS-FLI1. AHDC1 knockdown also reduced the transcriptional level of NR0B1 that harbors the GGAA microsatellite region within the promoter region. In addition, AHDC1 knockdown showed reduced cell growth in Ewing’s sarcoma cell lines but not non-Ewing’s cells. Together, we suggest that AHDC1 is one of the transcriptional co-regulators of EWS-ETS fusion proteins in Ewing’s sarcoma cells.

## Materials and Methods

### Cell culture

The A673 cell line was purchased from the European Collection of Authenticated Cell Cultures (ECACC) and cultured in Dulbecco’s Modified Eagle Medium (DMEM, Cat. No. 044-29765, Fujifilm-Wako chemical) supplemented with 10% heat-inactivated fetal bovine serum (FBS) and 1x Penicillin-Streptomycin Solution (Cat. No. 168-23191, Fujifilm-Wako chemical). The Seki cell line was established by Nojima *et al*. [18], purchased from the Cell Resource Center for Biomedical Research, Institute of Development, Aging and Cancer, Tohoku University (Cat. No. TKG 0725, Miyagi, Japan), and cultured in RPMI-1640 (Cat. No. 189-02025, Fujifilm-Wako chemical) with 10% FBS and 1x Penicillin-Streptomycin Solution. The NCR-EW2 cell line was cultured in RPMI-1640 with 10% FBS and 1x Penicillin-Streptomycin Solution. Human Embryonic Kidney cells 293 (HEK293) cells and hTERT RPE-1 (ATCC CRL-400) were cultured in DMEM with 10% FBS and 1x Penicillin-Streptomycin Solution. The Lenti-X™ 293T cell line was purchased from Takara-Bio (Cat. No. 632180) and cultured in DMEM with 10% FBS and 1x Penicillin-Streptomycin Solution. Seki, NCR-EW2, and Lenti-X293T cells were spread onto a 0.1% gelatin-coated dish.

### Plasmids

PrimeSTAR max polymerase (Cat. No. R045A, Takara-Bio) or KOD one polymerase (Cat. No. KMM-101, Toyobo) was used for precise cloning. The Welcome Sanger Institute kindly provided the pPB-LR5 [19] and pCMV-HyPBase [20] for the *piggyBac* system. The puromycin-resistant gene region, amplified from the linear puro marker (Cat. No. 631626, Takara-Bio), was inserted using the In-fusion HD cloning kit (Cat. No. 639648, Takara-bio) into the SpeI restriction site in the pPB-LR5, resulting in pPB-LR5-puro. The Tet3G-tet promoter-3xFLAG-EGFP fragment was amplified and inserted using the In-fusion HD cloning kit, resulting in the construction of pPBP-tet-3xEGFP. The BioID fragment was amplified from pcDNA3.1 mycBioID (Addgene: 35700) [10] and inserted into the pPBP-tet-3xEGFP after cutting at the KpnI and PmeI restriction enzyme sites, resulting in the construction of pPBP-tet-3xBioID-gs. The EWS-FLI1, EWS-ERG, and EWS-E1AF genes were amplified from pcDNA3-EWS-FLI1typeI, EWS-ERG, EWS-E1AF [21], and inserted into the PmeI restriction enzyme site of pPBP-tet-3xBioID-gs, resulting in pPBP-tet-3xBioID-EWS-FLI1, pPBP-tet-3xBioID-EWS-ERG, and pPBP-tet-3xEWS-E1AF, respectively. The AHDC1 gene (Genbank accession No. NM_001029882) was amplified from the cDNA of hTERT RPE-1 cells and inserted into the KpnI and PmeI restriction enzyme sites of pPBP-tet-3xEGFP, resulting in the construction of pPBP-tet-3xAHDC1. pGreenpuro shRNA cloning and expression lentivector was purchased from System Bioscience (Cat. No. SI505A-1, System Biosciences, LLC, CA). Primers for shRNA are shown in S1 Table. For shAHDC1, shFLI1, and shEWS, each primer shAHDC1-f and shAHDC1-r, shFLI1-f and shFLI1-r, shEWS-f and shEWS-r were annealed and inserted into the EcoRI and BamHI restriction enzyme sites of the pGreenpuro shRNA cloning vector. For measuring the transcriptional activity of EWS-FLI1, the NR0B1 promoter region was cloned from A673 genomic DNA, which was purified using a QIAamp DNA Mini Kit (Cat. No. 51304, QIAGEN), and inserted into the XhoI restriction enzyme site of pNL1.1[Nluc] vector (Cat. No. N1001, Promega), resulting in the construction of pNL1.1-NR0B1pro vector.

### Lentivirus production and transduction

For shRNA-expressing lentivirus production, 5 × 10^6^ Lenti-X 293T cells were cultured in 10 ml of DMEM medium on a plate coated with 0.1% gelatin for 24 h. Seventeen µg of pGreenpuro shRNA-expressing vector, ten µg of pCAG-HIVgp (RDB04394, RIKEN BRC) [22], and 10 µg of pCMV-VSV-G-RSV-Rev (RDB04393, RIKEN BRC) [22] were mixed with 111 µl of 1 mg/ml PEI MAX^®^ (pH7.5) (Cat. No. 24765-1, Polysciences) in Opti-MEM™ I Reduced Serum Medium (Cat. No. 31985070, Thermofisher Scientific) for 10 min. After changing the medium, the DNA mixture was treated and incubated for 6-24 h. The next day, after changing the medium, 100 µl of 500 mM sodium butyrate was added to enhance lentivirus production. The next day, 10 ml of medium were filtrated on 0.45 µm PVDF membrane of Millex-HV^®^ filter unit (Cat. No. SLHV R25 LS, MERCK KGaA), 3.5 ml of 4x PEG solution (32% PEG6000, 400 mM NaCl, 40 mM HEPES, adjusted to pH7.4) were added [23] for 1 h at 4°C and followed by centrifugation at 3000 rpm for 30 min at 4°C. The lentiviral pellet was mixed with 100 µl of PBS(-) containing 2.5% glycerol and stored at −80°C. Cells were cultured in a 12-well or 6-well plate for one day. The medium was replaced with a medium containing lentivirus particles and five µg/ml of DEAE-dextran to enhance lentivirus production [24] and incubated for two days. The medium was again cultured one more day for further analysis.

### Knockdown of target genes

Cells were cultured in a 6-well plate for a day; 100 pmol of siRNA was mixed with 4 µl of Lipofectamine™ RNAiMAX Transfection Reagent (Cat. No. 13778030, Thermofisher scientific) in Opti-MEM™ I Reduced Serum Medium and incubated for 10 min, followed by transfer to each well. A Stealth RNAi™ siRNA Negative Control Med GC Duplex #2 (siNC, Cat. No. 12935112, Thermofisher Scientific) was used as negative control siRNA. The AHDC1 siAHDC1 used was a Stealth RNAi™ siRNA (siRNA ID: HSS146954, Thermofisher Scientific).

### Reverse transcription-quantitative PCR (RT-qPCR)

Total RNA was purified using the FastGene^TM^ RNA basic kit (Cat. No. FG-80050, NIPPON Genetics). According to the procedure, cDNA was obtained using ReverTra Ace^®^ qPCR RT Master Mix with gDNA Remover (Cat. No. FSQ-301, Toyobo). qPCR was performed using Applied Biosystems™ PowerUp™ SYBR™ Green Master Mix (Cat. No. A25742, Thermofisher Scientific) with a StepOnePlus™ Real-Time PCR System (Thermofisher Scientific). The thermal cycling parameters followed PCR amplification conditions: 50°C for 2 min and 95°C for 2 min, 40 cycles of 95°C for 15 s, and 60°C for 1 min. The oligonucleotides used for RT-qPCR are shown in S2 Table. Relative quantification of each target was normalized by Glyceraldehyde-3-phosphate dehydrogenase (GAPDH). Error bars indicate the standard deviation of three independent biological replicates. Statistical analyses were performed by Student’s t-test.

### Western blot analysis

Cells were cultured and lysed in RIPA buffer [50 mM Tris-HCl pH8, 150 mM NaCl, 1% Nonidet P-40 (NP-40), 0.1% sodium dodecyl sulfate (SDS), 0.5% sodium deoxycholate, 10 µg/mL leupeptin, 10µg/mL aprotinin, 1mM Phenylmethylsulfonyl fluoride (PMSF), 1.5 mM Na_2_VO_4_, 10 mM NaF], sonicated for 10-15 s, and centrifugated at 15000 rpm for 15 min. Supernatants were used for the following procedure. According to the manufacturer protocol, protein concentration was determined by the Protein assay BCA kit (Cat. No. 297-73101, Fujifilm-Wako chemical). An equal amount of protein (10 µg) was applied in 5-20% SDS-polyacrylamide gel (SuperSep Ace; Cat. No. 199-15191, Fujifilm-Wako chemical) and transferred to the PVDF membrane (Immobilon-P; Millipore, Bedford, MA, USA). The membrane was blocked by 5% skimmed milk or 5% BSA for 1 h with shaking, incubated with a primary antibody at 4°C overnight, and a horseradish peroxidase (HRP)-conjugated secondary antibody for 1 h at room temperature with shaking. The membrane was visualized by Immunostar Zeta (Cat. No. 297-72403, Fujifilm-Wako chemical) and detected by using an Amersham Imager 600 (GE healthcare) or a WSE-6100 LuminoGraph I (ATTO Co., Ltd). Immunostaining for the PVDF membrane was performed using the following antibodies: FLI1 (1:1000 dilution, Cat. No. ab15289, Abcam), EWSR1 (1:2000 dilution, Cat. No. 11910S, Cell Signaling Technology), BRD4 (1:1000 dilution, Cat. No. AMAb90841, Sigma-Aldrich), DYKDDDDK (1:4000 dilution, Cat. No. 018-22381, Fujifilm-Wako chemical), NKX2-2 (1:1000 dilution, Cat. No. ab187375, Abcam), p27 Kip1 (D69C12) (1:2000 dilution, Cat. No. 3686, Cell Signaling Technology), GAPDH (D16H11) (1:5000 dilution, Cat. No. 5174, Cell Signaling Technology), BRG1 (A52) (1:2000 dilution, Cat. No. 3508, Cell Signaling Technology), AHDC1 (1:1000 dilution, Cat. No. HPA028648, Atlas antibodies), SOX2 (1:2000 dilution, Cat. No. GTX627405, GeneTex).

### Immunostaining

Cells were cultured and fixed using 4% Paraformaldehyde/PBS(-) for 15 min, permeabilized using 0.1% Triton X-100/PBS(-) for 15 min and blocked using 1% goat serum (Cat. No. 50062Z, Life technologies) for 15 min. Cells were incubated with primary antibodies at 4°C overnight and stained with secondary antibodies and five µg/ml 4’,6-Diamidino-2-phenylindole, dihydrochloride (DAPI). Primary antibodies used were DDDDK (1:1000 dilution, Cat. No. PM020, MBL), DYKDDDDK (1:2000 dilution, Fujifilm-Wako pure chemical), BRD4 (1:200 dilution, Sigma-Aldrich), or BRG1 (1:200 dilution, CST). Secondary antibodies used were Alexa Fluor 488-conjugated Goat anti-Mouse IgG (1:500 dilution, Cat. No. A-11001, Thermofisher Scientific), Alexa Fluor 488-conjugated Goat anti-Rabbit IgG (1:500 dilution, Cat. No. A-11034, Thermofisher Scientific), Alexa Fluor 555-conjugated Goat anti-Mouse IgG (1:500 dilution, Cat. No. A-21422, Thermofisher scientific), or Alexa Fluor 555-conjugated Goat anti-Rabbit IgG (1:500 dilution, Cat. No. A-21428, Thermofisher Scientific). SlowFade™ diamond antifade mountant (Cat. No. S36963, Thermofisher Scientific) was used as a mounting reagent. An All-in-One fluorescence microscope BZ-9000 (Keyence) was used for the observation in Fig 1. A Nikon A1R HD25 system confocal microscope with ECLIPSE Ti2E (Nikon) was used for the observation in Fig 5.

**Fig 1.**
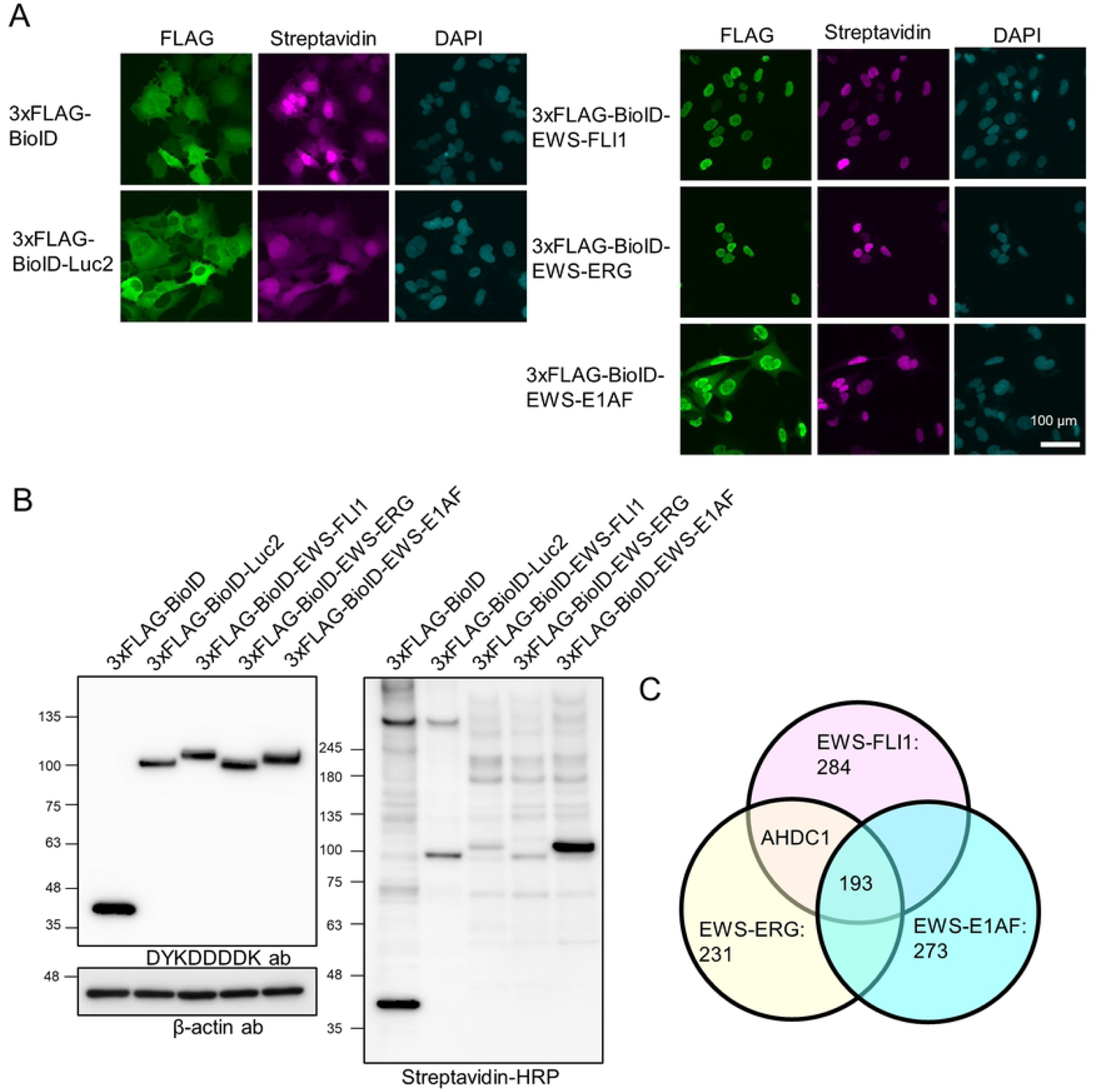
Identification of AHDC1 as a proximal protein of EWS-ETS fusion proteins. (A) 3xFLAG-BioID-tagged EGFP or EWS-ETS fusion proteins under the control of Tet-on promoter were expressed in A673 cells by 1 µg/ml doxycycline for 1 d. FLAG-tag or biotinylated proteins were stained with DYKDDDDK antibody or Alexafluor633-conjugated streptavidin, respectively, with DAPI. (B) Western blotting analysis of each BioID sample. FLAG-tag was stained with DYKDDDDK antibody. Biotinylated proteins were stained with streptavidin-HRP, and β-actin was stained as an internal control. (C) Identified protein numbers from each EWS-ETS fusion protein samples by mass spectrometry analysis. To determine whether AHDC1 is a proximal protein of EWS-ETS fusion proteins, we purified the biotinylated proteins again and detected AHDC1 (Fig 2A). The intensity of AHDC1 in the EWS-ETS protein sample was higher than in each BioID and BioID-Luc2 sample. Next, immunoprecipitation for AHDC1 was performed using FLAG-tagged AHDC1-expressing cells (Fig 2B). FLAG-tagged AHDC1 was immuno-precipitated with endogenous EWS-FLI1 protein compared to FLAG-tagged EGFP. FLAG-tagged EWS-FLI1 was also immunoprecipitated with endogenous AHDC1 compared to FLAG-tagged EGFP (Fig 2C). Moreover, endogenous EWS-FLI1 immunoprecipitants were included in AHDC1 with BRD4 and BRG1 (Fig 2D).

**Fig 2.**
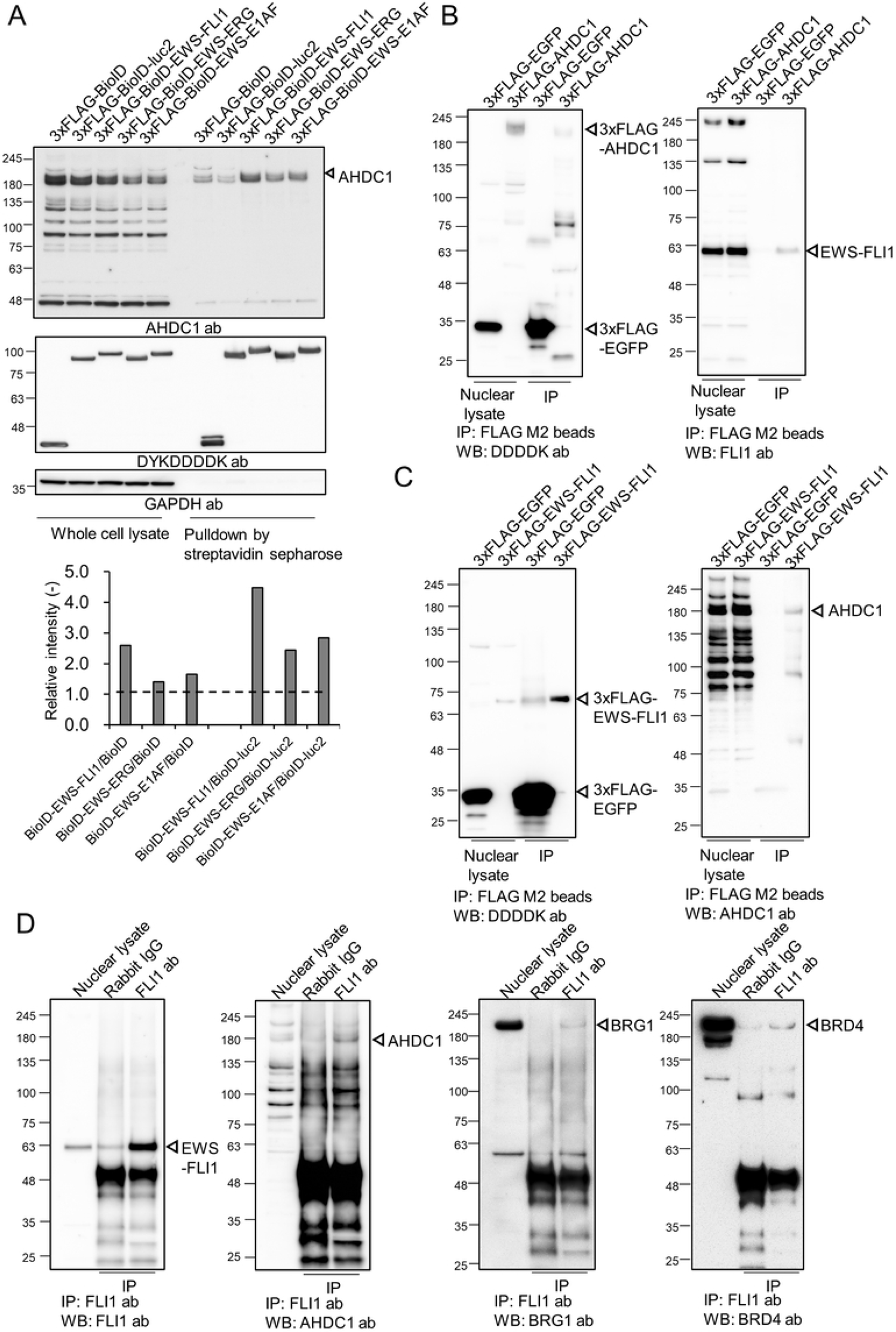
Immunoprecipitation of AHDC1. (A) Western blotting analysis after streptavidin-conjugated sepharose beads. Ten µg of proteins and one-tenth of pulldown input were used as a whole-cell lysate and a biotinylated protein sample, respectively. Band intensity was compared as a BioID or BioID-tagged Luc2. GAPDH antibody was used as a negative control. (B) Western blotting analysis of co-immunoprecipitated samples. 300 µg of nuclear lysate was mixed with FLAG M2 magnetic beads for immunoprecipitation. 5 µg of nuclear and one-fifth of the immunoprecipitation input were used for Western blotting. (C) Western blotting analysis of co-immunoprecipitated samples. 300 µg of nuclear lysate was mixed with FLAG M2 magnetic beads for immunoprecipitation. (D) Western blotting analysis of co-immunoprecipitated samples. 500 µg of nuclear lysate was mixed with FLI1 antibody and protein A/G magnetic beads for immunoprecipitation. 5 µg of nuclear lysate and one-fifth of the immunoprecipitation input were used for Western blotting.

### Biotin labeling in living cells and elution of the biotinylated proteins

Cells were induced to produce BioID fusion proteins and biotinylated BioID-proximal proteins by 1 µg/ml of doxycycline and 50 µM biotin for 24 h in a 10 cm dish. Isolation of the biotinylated proteins was followed by the Couzens *et al*. method [25]. After washing the cells with PBS 3 times, cells were lysed by 500 µl of RIPA buffer (50 mM Tris-HCl pH8, 150 mM NaCl, 1% NP-40, 0.1% SDS, 0.5% sodium deoxycholate, 1 mM EDTA, 1mM EGTA, 10 µg/ml leupeptin, 10 µg/ml aprotinin, 1 mM PMSF, 1.5 mM Na_2_VO_4_, 10 mM NaF). The cell lysate was incubated with 1 µl of Benzonase (Cat. No. 70746-3CN, Millipore), shaking on an icebox for 1 h, then sonicated for 15 s, and centrifuged at 15000 rpm for 15 min. The supernatant was mixed with 50 µl of streptavidin sepharose (Cat. No. 17-5113-01, GE healthcare), shaking at 4°C for 3 h after being washed with PBS once. After collecting beads by centrifugation, the beads were washed with RIPA buffer without protease inhibitors once, washed with TAP buffer (50 mM HEPES-KOH pH 8.0, 100 mM KCl, 10% glycerol, 2 mM EDTA, 0.1% NP-40) twice, and washed with 50 mM ammonium bicarbonate (pH 8.0) six times. The beads were incubated with 100 µl of 50 mM Tris-HCl (pH8.5) and 1 µl of 5 µg/µl dithiothreitol (DTT) with shaking at 37°C for 1 h. In addition, 1 µl of 12.5 mg/mL iodoacetamide was added to the beads and incubated with shaking at 37°C for 1h in the dark. The beads were finally added with 2.5 μl of 200 ng/µl Trypsin/Lys-C Mix (Cat. No. V5073, Promega) at 37°C with shaking overnight. The supernatant was collected by centrifuge, collected again after the beads were washed with 50 mM Tris-HCl (pH8.5), and added with 10 µl of 20% trifluoroacetic acid (TFA).

For the desalting step, styrene-divinylbenzene (SDB)-stage tip was washed with 20 µL of 0.1% TFA in 80% acetonitrile and further washed SDB-stage tip with 20 µl of 0.1% TFA in 2% acetonitrile. The peptide digest was transferred to the SDB-stage tip and trapped by centrifugation. The SDB-stage tip was washed with 20 µl of 0.1% TFA in 2% acetonitrile and 0.1% TFA in 80% acetonitrile. The peptides were eluted with 200 µl of 0.1% TFA, and 1-2 µl of peptide solution was applied for mass spec analysis.

### Liquid Chromatograph – Mass Spectrometry (LC-MS) analysis and label-free quantification

For BioID analysis, the peptide samples were subjected to a nano-flow reversed-phase (RP) LC-MS/MS system (EASY-nLC™ 1200 System coupled to an Orbitrap Fusion Tribrid Mass Spectrometer; Thermo Fisher Scientific, San Jose, CA) with a nanospray ion source in positive mode. Samples were loaded onto a 75-μm internal diameter × 2-cm length RP C18 precolumn (Thermo Scientific Dionex) and washed with loading solvent before switching the trap column in line with the separation column, a nano-HPLC C18 capillary column (0.075 × 125 mm, 3 mm) (Nikkyo Technos, Tokyo, Japan). A 60-min gradient with solvent B (0.1% Formic acids in 80% acetonitrile) of 5-40% for separation on the RP column equilibrated with solvent A (0.1% formic acid in water) was used at a flow rate of 300 nl/min. MS and MS/MS scan properties were as follows; Orbitrap MS resolution 120,000, MS scan range 350–1500, isolation window m/z 1.6, and MS/MS detection type was ion trap with a rapid scan rate.

All MS/MS spectral data were searched against entries for human in the Swiss-Prot database (v2017-06-07) with a mutant form of *E coli* biotin ligase (BirA) using the SEQUEST database search program using Proteome Discoverer 2.2 (PD2.2). The peptide and fragment mass tolerances were set to 10 ppm and 0.6 Da, respectively. For variable peptide modifications, oxidation of methionine and biotinylation of lysine, in addition to carbamidomethylation of cysteine for a fixed modification, were considered. Database search results were filtered by setting the peptide confidence value as high (FDR < 1%) for data dependent mass analysis data. For label-free quantification, the peptide and protein amount were calculated from the precursor ion intensities using the workflow of Precursor Ions Quantifier in PD2.2. The amount of mutant form of BirA quantified in each analysis was used for the bait normalization. ANOVA was performed to calculate the adjusted p-values to control experiments (BirA and BirA-Luc2) using the same workflow.

### Immunoprecipitation

Immunoprecipitation was performed with a slight modification of the following procedure [26]. Cells expressing 3xFLAG-tagged EGFP, ADHC1, or EWS-FLI1 under the control of a Tet-on system which was cultured in a medium containing 1 µg/ml doxycycline for 1 d, were washed by PBS(-) 3 times, collected in PBS(-) after scraping, and centrifuged at 450 *g* for 10 min at 4°C. The pellets were treated with 1 ml of hypotonic lysis buffer (10 mM KCl, 10 mM Tris pH 7.5, 1.5 mM MgCl_2_) supplemented with 1 mM DTT, 1 mM PMSF, 10 µg/ml Leupeptin, and 10 µg/ml Aprotinin for 15 min on ice followed by centrifugation at 400 *g* for 5 min at 4°C. Pellets were treated with 500 µl of hypotonic lysis buffer again and mixed by pipetting 10 times, followed by centrifugation at 10000 *g* for 20 min at 4°C. Pellets were treated with high-salt extraction buffer (0.42 M KCl, 10 mM Tris pH 7.5, 0.1 mM EDTA, 10% glycerol supplemented with 1 mM PMSF, 10 µg/ml Leupeptin, and 10 µg/ml Aprotinin) with 1 µl Benzonase (70746, Millipore), and gently shaken on an icebox for 30 min, followed by centrifugation at 20000 *g* for 5 min at 4°C. Supernatants were diluted by Milli-Q water to adjust to 150 mM salt concentration. 300 µg of nuclear lysate were topped up to 500 µl using IP wash buffer (150 mM KCl, 10 mM Tris pH 7.5, 0.1 mM EDTA, 10% glycerol) supplemented with 1 mM PMSF, 10 µg/ml Leupeptin, and 10 µg/ml Aprotinin. Fifty µl of Anti-FLAG magnetic beads (M8823, Millipore) were washed by PBS(-) once and rotated in 1 ml of 5% BSA/PBS(-) 1 h at 4°C. The nuclear lysate was mixed and rotated with anti-FLAG magnetic beads for 3 h at 4°C and washed using 1 ml of IP wash buffer 4 times. Beads were mixed with 50 µl of 2x SDS sample buffer at 95°C for 5 min. The supernatants were used for Western blotting analysis.

For endogenous protein immunoprecipitation, the nuclear lysate was collected using the above method. 500 µg of nuclear lysate was topped up to 500 µl using IP wash buffer and mixed with 10 µg of FLI1 [EPR4646] antibody (ab133485, Abcam) or rabbit normal IgG (Cat.148-09551, Wako pure chemical), followed by rotation at 4°C for 2 h. The nuclear lysate/IgG was mixed with 25 µl of Pierce™ Protein A/G Magnetic Beads (Cat. 88802, Thermofisher Scientific) with rotation at 4°C for 2h. The beads were washed with IP wash buffer 4 times and mixed with 50 µl of 2x SDS sample buffer at 95°C for 5 min. The supernatants were used for Western blotting analysis.

### Cell viability assay

Lentiviral-transduced cells were collected without a drug selection, and 1 × 10^3^ cells were spread in a 96-well plate. An equal volume of CellTiter-Glo^®^ 2.0 reagent (Cat. No. G924B, Promega) was transferred into each well and incubated for 5 min. After pipetting each well, the mixture was transferred into a 1.5-ml tube, mixed by a shaker for 10 min at room temperature, and luminescence was measured by a GloMax^®^ 20/20 Luminometer (Cat. No. E5311, Promega). To measure apoptotic activity, an equal volume of Caspase-Glo® 3/7 Assay System (Cat. No.G8090, Promega) was transferred into each well and incubated for 1 h and measured by a GloMax^®^ 20/20 Luminometer. *Spheroid formation assay.* Lentiviral-transduced cells were collected, and 1 × 10^4^ cells were spread in a PrimeSurface96U (Cat. No. MS-9096U, Sumitomo Bakelite). The medium was changed every 2 d, and photos were taken by an All-in-One fluorescence microscope BZ-810X (Keyence).

### Wound healing assay

Lentiviral-transduced cells were collected, 1 × 10^4^ cells were transferred into each culture-insert 2-well (Cat. No. ib80209, ibidi GmbH) and incubated overnight. After removing the culture-insert from the dish, cells were washed twice with PBS(-), transferred to DMEM medium without FBS, and pictures were taken by the BZ-810X microscope.

### Promoter reporter assay

1 × 10^4^ cells were cultured in a 96-well plate. The next day, 50 ng of the pNL1.1-NR0B1 vector was transfected with 0.1 µl of Lipofectamine™ Stem Transfection Reagent (Cat. No. STEM00003, Thermofisher Scientific) according to the procedure and incubated for 4 h. Three pmol of siNC or siAHDC1 stealth siRNA was incubated with 0.125 µl of Lipofectamine™ RNAiMAX Transfection Reagent (Cat. No. 13778030, Thermofisher scientific) in Opti-MEM™ I Reduced Serum Medium for 10 min and treated in each well. After 2 d, an equal volume of Nono-Glo Live-cell assay system (Cat. No. N2011, Promega) was added to each well and mixed by pipetting and shaking for 5 min and measured by a GloMax^®^ 20/20 Luminometer. Luminescence of no-transfected cells was subtracted from each sample. Error bars show the standard deviation of five independent biological replicates. Statistical analyses were performed by student’s t-test.

## Results

### Biotinylation of proximal proteins by BioID in A673 cells

For BioID-tagged EWS-ETS fusion protein expression, we constructed the *piggyBac* system under the control of the Tet-on system to regulate the gene expression. BioID-tagged EWS-FLI1, EWS-ERG, or EWS-E1AF-expressing plasmids were transfected into A673 cells with a hyperactive *piggyBac* transposase previously generated for applications in mammalian genetics [20]. After a puromycin selection, cells expressed each BioID-tagged gene by doxycycline with biotin. BioID alone or BioID-tagged Luc2 (firefly luciferase) were used as a negative control and labeled biotin to proximal proteins in all cell fractions (Fig 1A). In addition, BioID-tagged EWS-ETS fusion proteins were mainly localized in the nuclei. Next, we checked whether BioID-tagged EWS-ETS fusion proteins could biotinylate proximal proteins in A673 cells by Western blotting (Fig 1B). Streptavidin-HRP staining confirmed the appearance of various biotinylation bands.

We prepared three independent biological replicates for each cell line, collected biotinylated proteins by a streptavidin sepharose set up using the Couzens *et al*. method [25], and identified proteins by mass spectrometry analysis (Fig 1C and S3 Table). A total of 193 proteins were identified as proximal proteins shared by identified proteins list from the three fusion proteins (Abundance ratio: each fusion proteins list compared to BioID or BioID-Luc2 > 5, Abundance Ratio Adj. P-Value< 0.05). These common proteins list contained the chromatin remodeling complex (ARID1A, ARID2, BRG1, BCL11B, SMARCAL1, SMARCB1, SMARCC1, SMARCD1, and SMARCE1), splicing factors (SF1, SF3A1, SF3A2, SF3A3, SF3B2, SF3B4, and SCAF4), and super-enhancer-related proteins (BRD4, BICRA, MED11, MED13L, MED25, and MED30). AHDC1 was contained in the BioID-tagged EWS-FLI1 and EWS-ERG protein samples (Fig 1C). However, AHDC1 did not show a significant difference in the BioID-tagged EWS-E1AF protein list.

### AHDC1 knockdown affects gene expression of EWS-FLI1 target genes

To evaluate whether AHDC1 affects gene expression of EWS-FLI1, we treated A673 cells with siRNA for the AHDC1 knockdown experiment. AHDC1 knockdown showed reduced EWS-FLI1 protein expression level but not EWSR1 (Fig 3A). The nuclear receptor NR0B1 and the homeobox transcription factor NKX2-2 were up-regulated in Ewing’s sarcoma [27–29]. NR0B1 and NKX2-2 protein expression levels were reduced in siAHDC1-treated cells. Silencing of EWS-FLI1-bound GGAA microsatellite by a dCas9-KRAB system showed downregulation of NKX2-2 and SOX2 protein expression in A673 and SKNMC cells [30]. However, AHDC1 knockdown did not change the SOX2 protein level in A673 cells. We also tested whether AHDC1 knockdown reduces protein expression levels in other Ewing’s sarcoma cell lines. For this purpose, we treated Seki or NCR-EW2 cell lines, both of which have been established as Ewing’s sarcoma cells, with siAHDC1 RNA [18, 31]. EWS-FLI1 and NR0B1 were also downregulated in both cell lines (S1 Fig A and B). NKX2-2 was only downregulated in NCR-EW2 cells.

**Fig 3.**
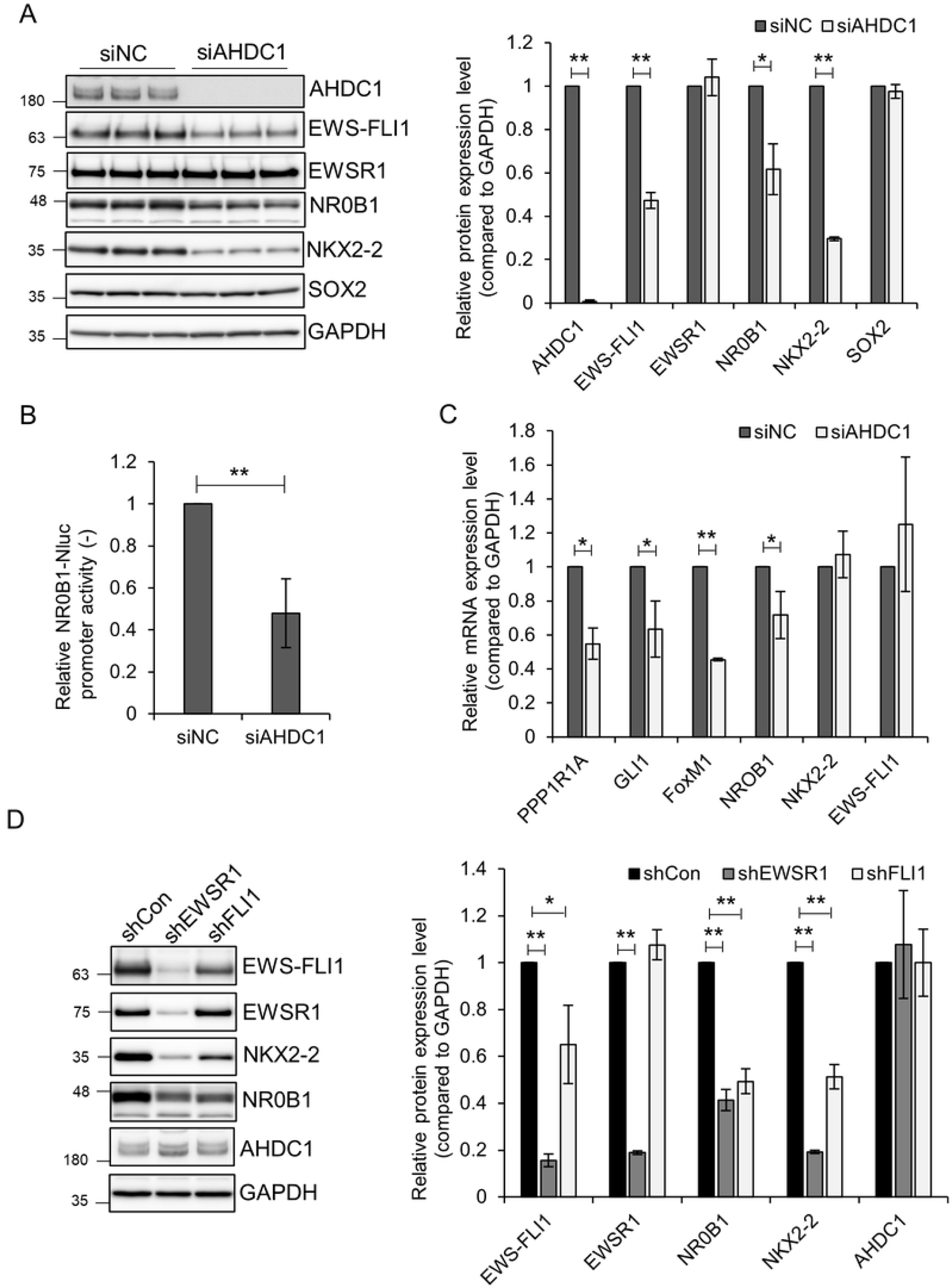
EWS-FLI1 knockdown reduces gene expression of EWS-FLI1 protein. (A) siAHDC1-treated A673 cells were cultured for 2 d. Each protein was detected by its respective antibody. (B) NR0B1 promoter-Nluc plasmid was transfected into A673 cells, incubated for 4 h, and treated with siRNA for 2 d. (C) siAHDC1-treated A673 cells were collected, total RNA was purified, and reverse-transcribed to cDNA. Each gene was quantified by the respective primer set using RT-qPCR. (D) Lentivirus expressing each shRNA was transduced into A673 cells for 3 d. GAPDH was used to normalize the relative values of Western blotting and RT-qPCR. Western blotting or RT-qPCR were quantified by three independent experiments. Nluc assay was quantified by five independent experiments. *P* values were calculated by the student’s t-test. * p<0.05; ** p<0.001.

The NR0B1 gene harbors EWS-FLI1-bound GGAA microsatellites within its own promoter region [32]. We cloned the NR0B1 promoter region upstream of Nanoluc and measured NR0B1 promoter activity in siAHDC1-treated cells (Fig 3B). AHDC1 knockdown showed downregulation of NR0B1 promoter activity in A673 cells. We also measured mRNA levels of target genes of EWS-FLI1 by RT-qPCR in siAHDC1-treated cells (Fig 3C). mRNA expression of PPP1R1A, GLI1, FoxM1, and NR0B1 genes highly expressed in Ewing’s sarcoma cells was dependent on EWS-FLI1 [32–35]. These genes were downregulated in siAHDC1-treated cells but not NKX2-2 and EWS-FLI1. To check whether EWS-FLI1 controls AHDC1 gene expression, EWS-FLI1 knockdown was performed (Fig 3D). AHDC1 protein expression level was not altered in shEWSR1 or shFLI1-treated cells. These results suggest that AHDC1 partially affects EWS-FLI1-mediated transcriptional activity but post-transcriptionally or post-translationally regulated EWS-FLI1 protein expression.

### AHDC1 knockdown attenuates cell growth in Ewing’s cells

EWS-ETS proteins are essential for the cell growth of Ewing’s sarcoma. To test whether AHDC1 affects cell growth in Ewing’s sarcoma cells, shAHDC1-expressing lentivirus was transduced in A673 cells (S2 Fig). EWS-FLI1, NR0B1, and NKX2-2 protein expression was reduced in shAHDC1-expressing cells as well as in siAHDC1-treated cells. After lentivirus transduction, cells were collected and spread again onto the 96-well microplate, resulting in the reduction of cell growth (Fig 4A). In addition, the spheroid culture of shAHDC1-expressing cells also showed reduced cell growth in a 3D-culture well (Fig 4B). Seki and NCR-EW2 cells were also treated with shAHDC1-expressing lentivirus, resulting in the reduction of cell growth (S3 Fig A). AHDC1 knockdown was also performed in HEK293 or hTERT RPE-1 cells as non-Ewing’s cell types (S3 Fig B). AHDC1 was expressed in both cell lines. NKX2-2 was weakly expressed in HEK293 cells, but shAHDC1 transduction did not alter the NKX2-2 protein expression level. In addition, HEK293 and hTERT RPE-1 cells did not show reduced cell growth after shAHDC1 transduction, suggesting that AHDC1 affects cell growth in Ewing’s sarcoma cells (S3 Fig C).

**Fig 4.**
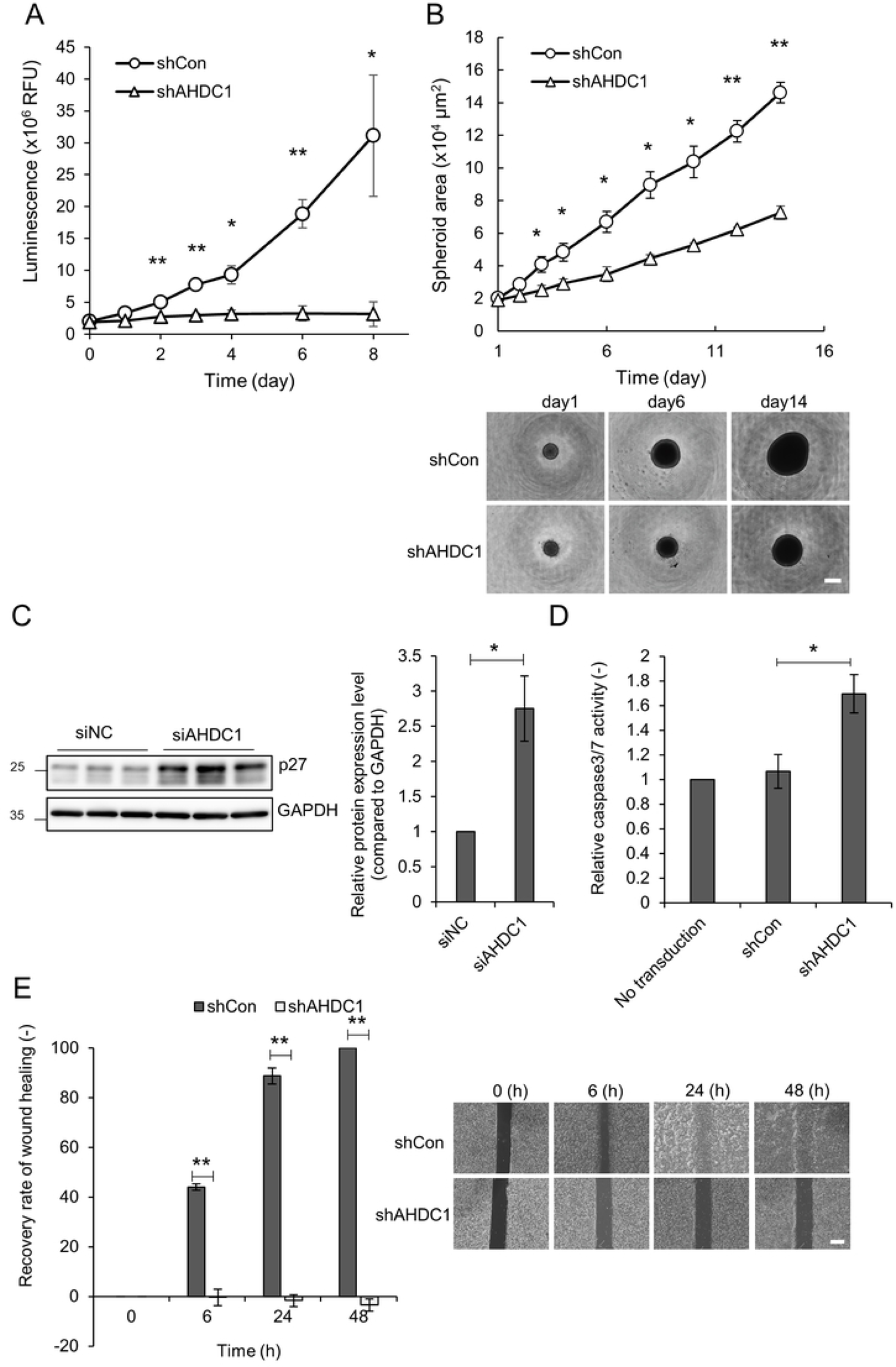
AHDC1 knockdown reduces cell growth in Ewing’s sarcoma cells. (A) Lentivirus expressing shRNA was transduced to A673 cells for 3 d. 1 × 10^3^ Cells were spread onto a 96-well plate and cultured again. Cell viability was determined by CellTiter-Glo2.0 on the indicated day. (B) shRNA-expressing cells were transferred into a 3D culture plate. The spheroid size was determined by a Keyence BZ-810X microscopy. Scale bar, 500 µm. (C) siAHDC1-treated A673 cells were cultured for 2 d. Western blotting analysis was performed by p27 and GAPDH antibodies. Relative values were normalized by GAPDH. (D) Lentivirus expressing shRNA was transduced to A673 cells for 3 d. Caspase activity was measured by a Caspase-Glo 3/7 Assay. (E) For the wound healing assay, 1 × 10^4^ shRNA-expressing cells were cultured for 1 d in a culture-insert, removed, and cultured again in a DMEM without FBS medium. Scale bar, 500 µm. *P* values were calculated by the student’s t-test. * p<0.05; ** p<0.001.

Next, we assessed cell cycle progression and apoptotic activity after AHDC1 knockdown. siAHDC1-treated cells presented an increased p27 protein level (Fig 4C). In addition, shAHDC1-expressing cells showed a high caspase activity level (Fig 4D). Finally, shAHDC1-expressing cells had reduced migration ability (Fig 4E). These results suggest that AHDC1 affects cell cycle progression, suppression of apoptosis, and reduced cell migration in Ewing’s sarcoma cells.

### AHDC1 knockdown reduces BRD4 and BRG1 protein expression

In our proximal proteins screening of EWS-ETS proteins, we also identified BRD4 and BRG1, both of which are super-enhancers and transcriptional regulators (S3 Table) [36]. BRD4 has been shown to interact with EWS-FLI1, and BRD4 inhibition by BET inhibitors results in reduced cell growth in Ewing’s sarcoma cells [7,8,37,38]. EWS-FLI1 recruited BRG1/BRM-associated factor (BAF) complexes containing BRG1 to the GGAA microsatellite region [39]. We tested whether BRD4 and BRG1 protein expression levels are affected by AHDC1 knockdown (Fig 5A). AHDC1 knockdown showed reduced BRD4 and BRG1 protein expression levels. Fluorescent protein-tagged AHDC1 localized in the nuclei in Hela cells [40]. We expressed FLAG-tagged AHDC1 by using the *piggyBac* system under the control of the Tet-on system in A673 cells and stained with BRD4 and BRG1 (Fig 5B). AHDC1 was co-localized with endogenous BRG4 but not BRG1. Thus, AHDC1 may regulate not only EWS-FLI1 but also BRD4 protein expression level in Ewing’s sarcoma cells.

**Fig 5.**
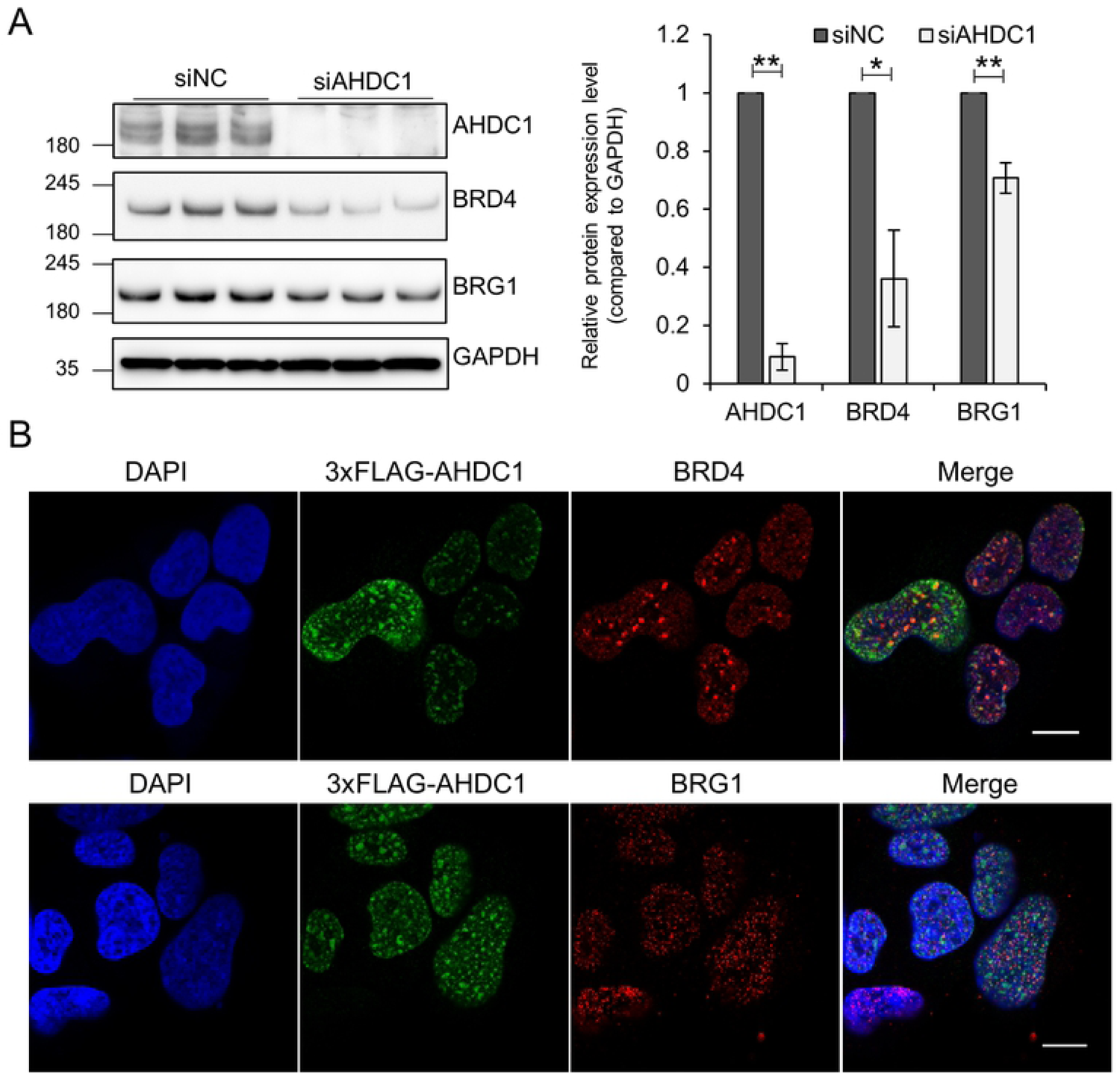
AHDC1 knockdown reduces BRD4 and BRG1 in A673 cells. (A) siAHDC1-treated cells were collected for Western blotting. Each antibody detected the respective protein, and the relative value was normalized by GAPDH. (B) 3xFLAG-AHDC1 was induced by 0.1 µg/ml doxycycline for 1 day, fixed, and permeabilized. Scale bar, 10 µm. *P* values were calculated by the student’s t-test. * p<0.05; ** p<0.001.

## Discussion

Proximal protein identification using new tools such as APEX2, BioID, or their derivatives has been a promising tool for biochemical approaches in vitro or in vivo [9]. In this study, we isolated AHDC1 as a proximal protein of the EWS-ETS proteins using the screening of the BioID system. AHDC1 was necessary to grow Ewing’s sarcoma cells but not non-Ewing’s sarcoma cells such as HEK293 or hTERT RPE-1 cells. In addition, AHDC1 affected gene expression of EWS-FLI1 target genes. Thus, AHDC1 may be one of the regulators for oncogenic function in Ewing’s sarcoma cells.

The Xia-Gibbs syndrome has been identified as a de novo heterozygous truncating mutation of AHDC1 [12]. To date, more than 100 cases of mutations related to the diagnosis of the Xia-Gibbs syndrome have been reported [13]. Not only heterozygous mutations of AHDC1 but also micro-duplication of the genome containing the AHDC1-coding region showed similar symptoms [41]. Thus, deregulation of AHDC1 gene expression affects the developmental process. AHDC1 has an AT-hook DNA binding motif, a PDZ motif, and other conserved domains within the coding sequence [40]. Feng *et al*. showed that AHDC1 expression was highly expressed in cervical cancer cells compared with immortalized cervical epithelium, and its expression was regulated by a long noncoding RNA, LINC01133 [42]. However, the molecular mechanisms for AHDC1 in cancer cells are still unclear.

EWSR1 is an RNA-binding protein comprising FET family proteins (FUS, TAF15, and EWSR1). EWSR1 is also one of the paraspeckle components that is a subcellular body in the nucleus and co-localized with SFPQ1, NONO, and PSPC1 [43, 44]. AHDC1 was also isolated as one of the paraspeckle components co-localized with EWSR1 [43]. Khayat *et al*. showed that wild-type AHDC1 localized in the nucleus, and Xia-Gibbs patients with mutation of AHDC1 have disrupted wild-type AHDC1 localization in HeLa cells [40]. Our proximal proteins screening of EWS-ETS fusion proteins did not isolate SFPQ, NONO, or PSPC1. However, CPSF5 (NUDT21), CPSF6, and CPSF7 that were isolated as paraspeckle components and are the components of the cleavage factor Im (CFIm) complex that brings about cleavage of 3’UTR of mRNA for polyadenylation were isolated as proximal proteins of EWS-ETS fusion proteins (S3 Table) [43]. These results suggest that some paraspeckle components may interact with transcriptional complexes with EWS-ETS fusion proteins.

FET family proteins comprising FUS, EWSR1, and TAF15 are not only involved in neurodegenerative disease but also act as oncoproteins in sarcoma or leukemia by chromosomal translocation. The N-terminal region of FET family proteins comprising SYGQ-rich regions interacts with the SWI/SNF chromatin remodeling complex containing BRG1 [39, 45]. In our screening, the chromatin remodeling complex containing BRG1, ARID1A, SMARCC1, SMARCD1, SMARCE1, SMARCB1, and SMARCAL1 were isolated as proximal proteins of EWS-ETS fusion proteins (S3 Table). EWS-FLI1 recruits BRG1 to open the chromatin structure at the GGAA microsatellite region [39]. In our observations, AHDC1 contributed to maintaining BRG1 protein expression level but did not co-localize with BRG1 (Fig. 4A and B). We postulate that the SWI/SNF chromatin remodeling complex may regulate EWS-FLI1 transcriptional activity with AHDC1.

AHDC1 did not regulate the gene expression of EWS-FLI1 at the transcriptional level (Fig. 3A and C). This means that AHDC1 might affect the protein stability of EWS-FLI1 at the post-translational level or post-transcriptional level. EWS-FLI1 is controlled to be degraded by the proteasomal machinery through a single lysine ubiquitination site [46]. AHDC1 might stabilize super-enhancers containing BRD4. In addition, FLAG-tagged AHDC1 expression co-localized with BRD4 in Ewing’s sarcoma cells (Fig. 5B). We hypothesize that AHDC1 might be one of the accessory proteins needed to stabilize super-enhancers containing EWS-FLI1 and BRD4 in Ewing’s sarcoma cells.

## Conflicts of Interest

The authors declare no conflicts of interest concerning this study.

## Acknowledgments

pcDNA3.1 mycBioID was a gift from Kyle Roux (Addgene plasmid # 35700; http://n2t.net/addgene:35700; RRID:Addgene_35700). The RIKEN BRC provided pCMV-HIVgp and pCMV-VSV-G-RSV-Rev through the National BioResource Project of the MEXT/AMED, Japan. The authors would like to acknowledge the technical support of the Advanced Research Promotion Center, Health Sciences University of Hokkaido. The authors would like to acknowledge the technical expertise of the DNA core facility of the Center for Gene Research, Yamaguchi University, supported by a grant-in-aid from the Ministry of Education, Science, Sports, and Culture of Japan. The authors would also like to acknowledge contributions to the proteome analyses of the staff members of the Proteomic Analysis Core System at the General Research Core Laboratory, Kumamoto University Medical School. We also thanks to Jun-ichi Hamada for a technical assistance.

## Author Contributions

TK designed performed experiments and wrote the paper. DK and NA performed mass spectrometry analysis, analyzed the raw data, and wrote the paper. BB and YK wrote the paper and discussed about the experimental concept. OH, TM, DP, RT, TO, YA, MT, MFS, KN, and MK gave technical support and conceptual advice.

## Supporting information

**S1 Fig. AHDC1 knockdown of Seki and NCR-EW2 cells.** A. siAHDC1-treated Seki cells were cultured for 2 d. Each protein was detected by its respective antibody. B. siAHDC1-treated NCR-EW2 cells were cultured for 2 d. Each protein was detected by its respective antibody. *P* values were calculated by the student’s t-test. * p<0.05; ** p<0.001.

**S2 Fig. AHDC1 knockdown of A673 cells by lentivirus expressing shRNA.** Lentivirus expressing shRNA was transduced to A673 cells for 3 d. Each protein was detected by its respective antibody. *P* values were calculated by the student’s t-test. * p<0.05; ** p<0.001.

**S3 Fig. AHDC1 knockdown shows reduced growth of Seki and NCR-EW2 by lentivirus expressing shRNA.** Lentivirus expressing shRNA was transduced to Seki or NCR-EW2 cells for 3 d. 1 × 10^3^ Cells were spread onto a 96-well plate and cultured again. Cell viability was determined by CellTiter-Glo2.0 on the indicated day. B. Lentivirus expressing shRNA was transduced to HEK293 or hTERT RPE-1 cells. Each protein was detected by its relative antibody. C. Lentivirus expressing shRNA was transduced to HEK293 or hTERT RPE-1 cells for 3 d. 1 × 10^3^ Cells were spread onto a 96-well plate and cultured again. Cell viability was determined by CellTiter-Glo2.0 on the indicated day. *P* values were calculated by the student’s t-test. * p<0.05; ** p<0.001.

**S1 Table. Primers for shRNA.**

**S2 Table. Primers for RT-qPCR.**

**S3 Table. LC-MS data of each BioID samples.**

**S1 Raw images**

## Notes

### Competing Interest Statement

The authors have declared no competing interest.

